# Structural Dynamics and Catalytic Mechanism of ATP13A2 (PARK9) from Simulations

**DOI:** 10.1101/2021.06.01.446648

**Authors:** Teodora Mateeva, Marco Klähn, Edina Rosta

## Abstract

ATP13A2 is a gene encoding a protein of the P5B subfamily of ATPases and is a PARK gene. Molecular defects of the gene are mainly associated with variations of Parkinson’s Disease (PD). Despite the established importance of the protein in regulating neuronal integrity, the three-dimensional structure of the protein currently remains unresolved crystallographically. We have modelled the structure and reactivity of the full-length protein in its E1-ATP state. Using Molecular Dynamics (MD), Quantum cluster and Quantum Mechanical/Molecular mechanical (QM/MM) methods, we aimed at describing the main catalytic reaction, leading to the phosphorylation of Asp513. Our MD simulations suggest that two positively charged Mg^2+^ cations are present at the active site during the catalytic reaction, stabilizing a specific triphosphate binding mode. Using QM/MM calculations, we subsequently calculated the reaction profiles for the phosphoryl transfer step in the presence of one and two Mg^2+^ cations. The calculated barrier heights in both cases are found to be ∼12.5 and 7.0 kcal mol^-1^, respectively. We elucidated details of the catalytically competent ATP conformation and the binding mode of the second Mg^2+^ cofactor. We also examined the role of the conserved Arg686 and Lys859 catalytic residues. We observed that by lowering significantly the barrier height of the ATP cleavage reaction, Arg686 had significant effect on the reaction. The removal of Arg686 increased the barrier height for the ATP cleavage by more than 5.0 kcal mol^-1^ while the removal of key electrostatic interactions created by Lys859 to the γ-phosphate and Asp513 destabilizes the reactant state. When missense mutations occur in close proximity to an active site residue, they can interfere with the barrier height of the reaction, which can halt the normal enzymatic rate of the protein. We also found large binding pockets in the full-length structure, including a transmembrane domain pocket, which is likely where ATP13A2 cargo binds.

## Introduction

The ATP13A2 gene has emerged as one of the genes strongly correlated with Parkinson’s disease (PD) and is also known as PARK 9^1,2^. The ATP13A2 gene encodes the P5B ATPase ATP13A2 which has attracted interest as an enzyme implicated in a range of neurodegenerative disorders: Spastic Paraplegia (SPG78), Kufor-Rakeb syndrome, neuronal ceroid lipofuscinosis and various other types of neurodegenerative disorders^2–6^. Currently, a multitude of molecular defects associated with the gene have been identified,^2,4,7–10^ including loss-of-function missense mutations of the protein^4,5,7–16^. The precise role of most missense mutations remains unexplored, as well as the overall structural dynamics of the protein.

ATP13A2 belongs to the haloacid dehydrogenase-like (HAD) superfamily of enzymes which all share a hydrolase fold. The HAD superfamily is very diverse, encompassing phosphoesterases, P-type ATPases, phosphonatases, dehalogenases, and sugar phosphomutases, which act on a wide range of substrates, typically catalyzing carbon or phosphoryl group transfer reactions^17^. Phosphotransferase enzymes typically require Mg^2+^ cofactor for their catalytic activity^18,19^. ATP13A2, in particular, belongs to the big family of P-type ATPases which is split in five distinct subfamilies: P1, P2, P3, P4, P5^6^. Most of these proteins are well studied and have resolved crystallographic structures, including ones in different functional states. ATP13A2 is part of the least studied subfamily P5B, which remains the only subfamily without any three-dimensional structures resolved.

The cytoplasmic domains of the protein include: Nucleotide-binding domain N, Phosphorylation domain P and an actuator domain A (Fig. 1A). Additionally, a transmembrane domain (T) connects the catalytic domains located in the cytoplasm to the extracytoplasmic area (Fig. 1B). Interestingly, the various missense mutations of the ATP13A2 protein currently described in the literature^5,8,9,11,14^ are not confined to one spatial region (Fig. 1A), but are scattered across the entirety of the protein and encompass all domains. The catalytically active domains, N, P and A are involved in ATP binding, ATP cleavage, auto and de-phosphorylation. ATP13A2 has been classified as a membrane transporter protein^6^ with the proposed candidates ranging from: heavy metals^2^, to Ca^2+^ cations^20^ and polyamine spermidine (SPD)^21,22^. Recent studies have revealed the role of ATP13A2 in polyamine export^23^. All enzymes belonging to the HAD superfamily contain a specific form of the Rossmannoid fold. This fold has two characteristic features that distinguish it from other superfamilies with Rossmannoid type-folds: a β-hairpin motif (also called a “flap”) located immediately downstream of the first β-strand of the core Rossmanoid fold and a single helical turn (“the squiggle”)^17^. This is important for ATP13A2 and other P-type ATPases, as these motifs provide mobility which allows the protein to alternate between the E1 “open” conformation (before the binding of any cargo) and the E2 “closed” conformation. The E1 state is associated with the binding and subsequent cleavage of ATP. In this state, the protein has a high affinity for the cargo that is to be transported from the cytoplasm to the other side of the membrane. In ATP13A2, the ATP cleavage reaction results in autophosphorylation of the strictly conserved Asp513. Similarly, the E2 state is associated with the process of dephosphorylation of the aspartate^6^. In this work, we are interested in the change from the E1-ATP to the E1P functional state.

**Figure 1.**
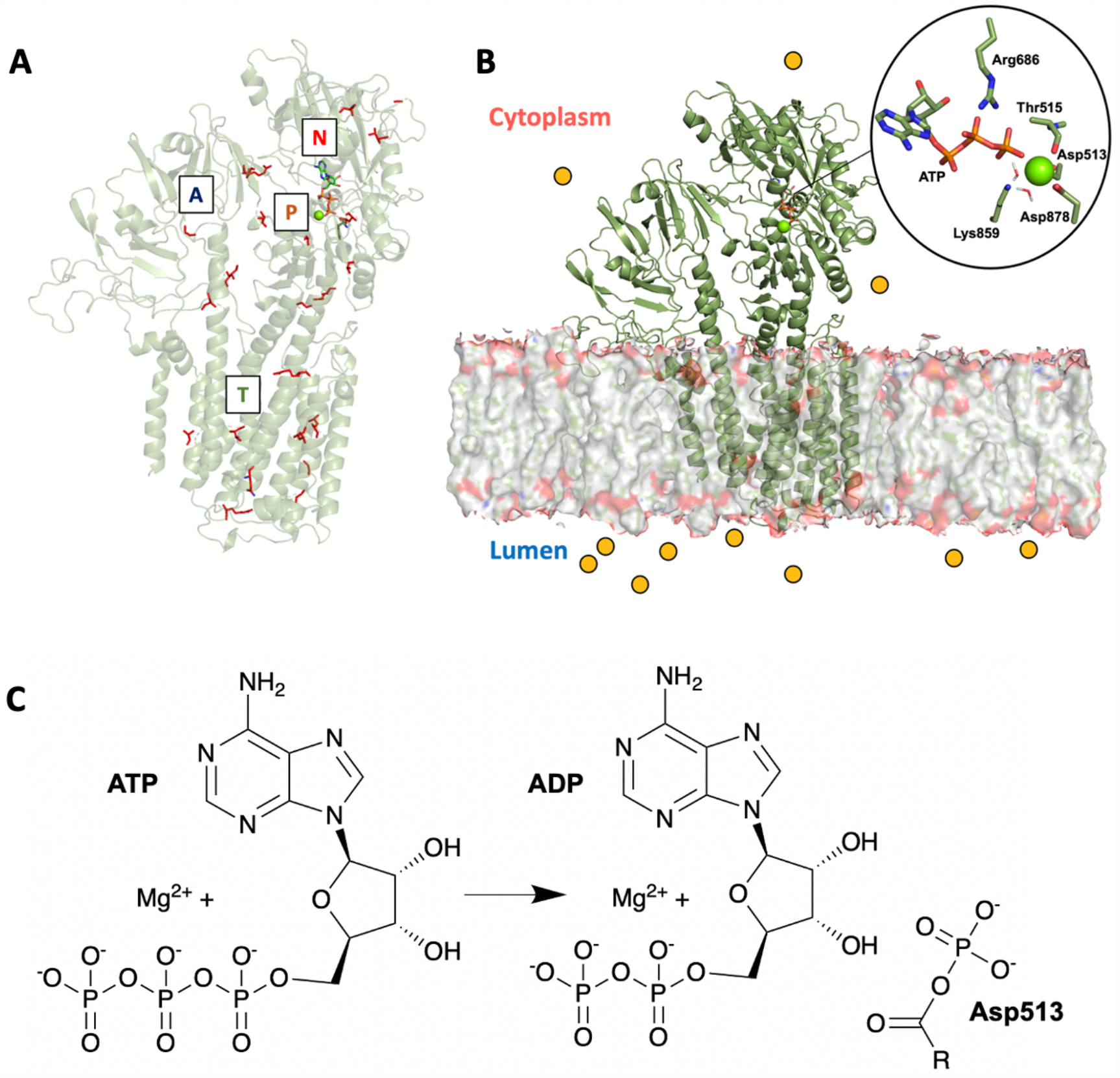
**(A)** Three-dimensional homology model of the ATP13A2 protein depicting missense mutations identified clinically^10,13,14,24,25^ (red sticks) on the protein (green cartoon) and **(B)** a homology model of ATP13A2 with the transmembrane domain buried in the lipid-rich membrane while the N, P and A domains are located in the cytoplasm of the cell. **(C)** Substrate in the E1 state and the products of the autophosphorylation reaction.

The active site motif DKTGT is strictly conserved among all P-type ATPases^26^. Two other amino acids, which are highly conserved and located immediately in the active site, are Arg686 and Lys859. Arg686 and Lys859 were structurally conserved in all enzymes whose crystal structures were used further in this study^27–32^, Fig. S1.

Currently, none of the proteins within the P5B ATPase family have been resolved crystallographically, including ATP13A2, therefore, no three-dimensional structure is resolved in any of the functional states of the protein. Nevertheless, ATPases of the P2A and P2C subfamilies, which are highly homologous, are available with experimentally determined structures. Most recently, a crystal structure of the P5A ATPase ATP13A1 was resolved, which is currently the most homologous protein to ATP13A2 whose three-dimensional structure has been determined^33^. Many of the three-dimensional P-type ATPase structures contain active-site bound ATP-analogues, typically with synthetic non-hydrolysable derivatives of ATP, such as ACP or AMPPCP^27–29^. Most of these structures feature only one bound active site Mg^2+^ cation^27–29^. However, there are structures obtained with ADP and AlF_3_ that feature two Mg^2+^ cations^30,32,34^ bound in the active site. This brings up the question whether one or two Mg^2+^ ions are present and/or required for the phosphoryl transfer to proceed? Importantly, due to the different charge distribution of the synthetic derivative analogues, the second ion coordination may be captured incorrectly or not at all. There are no structures which have been crystallised with the catalytically competent ATP, and, it is currently unknown what the precise ATP conformation during the phosphoryl transfer reaction is, especially in terms of its proposed second Mg^2+^ coordination^34^. This leaves open the question of what the precise ATP-Mg^2+^ coordination is, as well as overall Mg^2+^-Mg^2+^ distance and position.

Currently, the catalytic mechanism is not available using atomistic details for any P5 ATPase. Previous short Molecular Dynamics (MD) simulations were carried out on the P2A Endoplasmic Reticulum Ca^2+^-ATPase^35^, however, those did not provide any detailed insight on the catalytic mechanism or overall conformational dynamics of the protein. Multiscale reactive molecular dynamics (MS-RMD) and free energy sampling have been used to quantify the free energy profile and timescale of the proton transport in SERCA^36^. Quantum Mechanical/Molecular Mechanical (QM/MM) calculations have also previously been performed for the Phosphoserine Phosphatase, which also belongs to the large HAD-like superfamily of proteins, but it is classified in a distinctly different family^37^. We therefore performed MD, QM cluster and QM/MM calculations with the aim to describe this important catalytic mechanism and quantify the role of active site residues and active site cations in the phosphoryl transfer reaction in the catalytically competent E1-ATP functional state. Our QM/MM calculations found that Arg686 had a significant effect on the barrier height similarly to arginine fingers, however interacting with the β-phosphate of the ATP backbone^38^. Accordingly, experimental data from mutagenesis studies of the Ca^2+^-ATPase suggests that this conserved arginine is detrimental to the ATPase activity of the homologous enzyme^30^. We further show the precise effect on the barrier height of Lys859, which is similarly very important through interactions with both the γ-phosphate of the ATP and Asp513 in the reactant state. The results presented in this work can suggest how missense mutations disrupting crucial barrier-lowering interactions can have abolishing effect on the enzymatic activity of ATP13A2 and other P5B ATPases with homologous active site, specifically G877R ^5^.

## Methods

### Homology Modelling

The E1-ATP-bound state was modelled based on the Endoplasmic Reticulum Ca^2+^-ATPase (SERCA), (PDB code: 3tlm)^29^. The position of the ATP molecule and the Mg^2+^ cation was based on the position of the ACP moiety and the Mg^2+^ cation, respectively, in this crystal structure^29^. The crystal structure has the position of only one Mg^2+^ resolved, hence, the initial model contained only one Mg^2+^. The modelling server used was SWISS-MODEL^39^ and the sequence of ATP13A2 was obtained from Uniprot^40^ for Homo Sapiens (Uniprot code: Q9HD20). To account for the most recent crystal structure available of ATP13A1, ATP13A2 was also modelled based on the P5A ATPase ATP13A1 (PDB code: 6xms)^33^. Both templates result in the same three-dimensional structure in the active site region of interest (Fig. 2).

**Figure 2.**
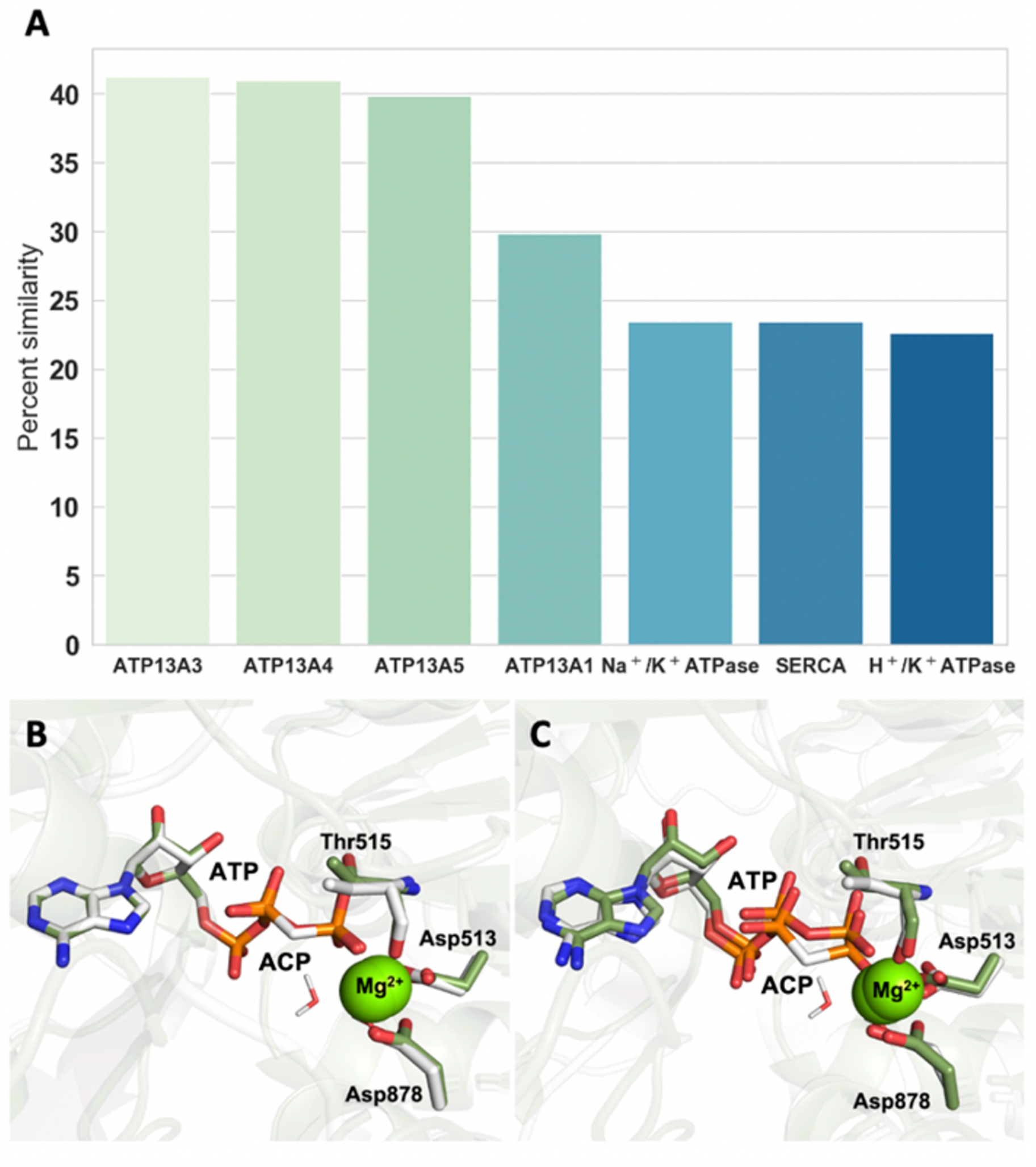
**(A)** Most similar human proteins to ATP13A2 based on overall fold, percent of sequence similarity and overall query coverage, ordered from most to least similar. **(B)** Active site of the Endoplasmic Reticulum Ca^2+^-ATPase (SERCA) in E1 state with bound ACP molecule and 1 Mg^2+^ cation, (PDB code: 3tlm^29^, grey) and our homology model of ATP13A2, (green sticks). **(C)** Active site of ATP13A1 in the E1 state with bound ACP molecule and 1 Mg^2+^ cation, (PDB code: 6xmq, grey sticks) and our homology model of ATP13A2, (green sticks).

### Molecular Dynamics (MD) Simulations

All Molecular Dynamics simulations were performed by using the program NAMD ^41^. The force field used in the simulations was CHARMM36^42^ with periodic boundary conditions and to evaluate the non-bonded long-range interactions the particle mesh Ewald method^43^ was utilised with a 12 Å cut-off. The NPT ensemble was maintained with a Langevin thermostat (303 K) and an anisotropic Langevin piston barostat (1 atm). The simulation was repeated at 309.15 K. The box type was rectangular, 22.5 Å per side. The water model was TIP3P^44^. To neutralize the system 0.15 M KCl solution was added. The ion placing method was by distance. The energy of the system was minimized via steepest descent algorithm, followed by a standard six-step equilibration for membrane-embedded systems with restrained heavy atoms via standard CHARMM-GUI^45^ procedure with a time step of 2 fs. The first step of the equilibration was done with a time step of 1 fs. SHAKE algorithm^46^ was deployed to constraint the covalent bonds involving hydrogen atoms. The equilibration was followed by 100 ns production with all the atoms completely unconstrained and free to move. A second 100 ns MD simulation was performed where the coordinates of the ATP molecule were fixed in their original position (following the crystal structure coordinates of the homologous template) in order to preserve the original coordinates of the crystal structure ACP (PDB code: 3tlm)^32^. The protein, solvent and ions were completely unconstrained. The PPM server was used for orientation of the protein in the membrane^47^. The membrane had the following composition: 40% Cholesterol, 30% Phosphatidylcholine lipids (PC) and 30% Phosphatidylethanolamine lipids (PE) in order to mimic a membrane environment in a lysosome-like cell. The same protocol was repeated for the second homology model of ATP13A2, which was based on the P5A ATPase ATP13A1.

### QM Cluster Calculations

All of the QM cluster calculations were performed by using the Gaussian 09 program^48^. The QM region was treated with the B3LYP hybrid density functional^49^ and the 6-31+G* basis set^50^. The QM region consisted of 2 Mg^2+^ cations, 6 water molecules, the side chain of Asp513, the side chain of Asp878 and Thr515, as well as the full ATP molecule (Fig. S2). The geometry optimization followed standard QM cluster procedure where the C atom where the amino acids are truncated, is frozen. Where a single C-C bond is cut, three H atoms are added to satisfy the C atom valency. The exact atoms which are frozen and truncated in the calculation are illustrated in Fig. S2. The second Mg^2+^ cation in the starting geometry of G1 (Fig. S3A) was placed based on alignment with the crystal structure of the Na^+^/K^+^ -transporting ATPase, which has full sequence conservation within 4.5 Å of the Mg^2+^ ions and is resolved with ADP and two Mg^2+^ cations (PDB code: 3wgu)^32^. The starting structure of G3 was taken from a snapshot of the last nanosecond of the unconstrained MD simulation (Fig. S3C). In G2 and G4 the second Mg^2+^ ion was placed by a manual initial guess.

### Quantum Mechanical/Molecular Mechanical Methods (QM/MM)

All QM/MM calculations were performed by Q-Chem^51^, coupled with CHARMM^52^. The QM region contained: the Mg^2+^ cation/s, six water molecules, the phosphate chain of ATP, the side chain of Asp513, the side chain of Asp878, Thr515, the side chain of Lys859, Arg686, Gly516 and the main chain of Lys514. The QM region during the RCS was treated with the B3LYP hybrid density functional and 6-31+G* basis set^49,50^. The MM region contained all residues and solvent within 25 Å of the QM region. The residues included in the QM region were separated from the rest of the chain by cleaving homonuclear C-C bonds and introducing link atoms, which were treated as hydrogen atoms in the QM calculations. An initial energy minimization was carried out, which constituted 1000 steps via the SCF DIIS algorithm. Each minimized geometry was supplied for the RCS as a reactant state starting point. The active site containing two Mg^2+^ ions was obtained by aligning the ATP and second Mg^2+^ ion from the QM cluster optimization and translating the optimized geometry to the QM/MM model. The conformation of the ATP molecule and the position of the Mg^2+^ ion in the one Mg^2+^-model were taken directly from the one observed in the crystal structure of the E1-ATP state of SERCA^29^, however, substituting one carbon atom of the crystal structure ACP molecule to a phosphorus, in order to have the catalytically-active ATP. This structure was further minimized for 1000 steps. We defined the reaction coordinate by the distance from the nucleophile to the phosphorus O_2D_-P_G_ (R1) and the phosphorus and the leaving group P_G_-O_3B_ (R2). Starting from the reactant state and moving along this coordinate, we simultaneously decrease the R1 distance and increase the R2 distance, to reach the product state. The distances were changed linearly. This forward-backward scanning is performed until energy convergence is observed. The solvent molecules in the QM region do not undergo reorganization from reactant to product state so the system has not been constrained additionally. 40 minimization steps were completed each time before a datapoint was recorded during the RCS. All presented scans are converged and show the forward scan direction, going from the reactant to the product state. Six systems were independently minimized. The minimized structure of each was supplied for a starting structure (reactant) of the RCS. For the purpose of this manuscript, “all amino acids” refers to Asp513, Asp878, Thr515, Lys859, Arg686, Gly516 and Lys514. Each system studied and its corresponding QM-region overall charge and atoms is summarised in Table S1.

### Pocket Analysis

Twenty frames were extracted from the 100 ns MD trajectory of the unconstrained simulation. The frames were spaced equidistantly and covered the duration of the MD simulation (Table S2). The twenty biggest pockets were calculated for each frame, using the Pymol^53^ plugin PyVOL^54^. To find pockets on the surface of ATP13A2, PyVOL^54^ was provided with the protein chain only without the ATP or any of the Mg^2+^ ions. The four biggest pockets in terms of surface area were chosen for further analysis. Upon inspection, it was observed that the biggest pockets for every frame were observed in the N-binding domain where ATP binding normally occurs, and in the transmembrane domain, respectively. For this work, we define a pocket as an “ATP-binding pocket” if the pocket was found in a location within 1.5 Å of the ATP in the respective frame from the simulation. We define a “transmembrane pocket” if a pocket is located within 1.5 Å of where small inorganic ion binding has been observed in the crystal structures^29,55^ of homologous proteins. If a pocket was calculated by PyVOL^54^ to be occupying this area of interest, it was recorded as found. If it was not calculated within this area, it was recorded as not found (Table S2). In this way, we were able to calculate the frequency of occurrence of those two pockets (Table S2). Other pockets which were found consistently are illustrated in S4. However, their occurrence was not as consistent as for the ATP pocket and the main transmembrane pocket. The results presented are obtained from the MD simulation of the homology model based on SERCA, but the same analysis was performed on a homology model based on ATP13A1 and the main transmembrane pocket was consistently found as well.

## Results

### Homology Modelling

To identify the closest protein sequences to ATP13A2 and any available structural information on these, we performed a BLAST^56^ and FASTA^57^ searches. The most similar proteins in humans, by sequence similarity, are as follows: ATP13A3, ATP13A4, ATP13A5 (Fig. 2A), which are all part of the P5B ATPase protein subfamily. As it was proposed earlier^23^, those proteins have high conservation in the substrate binding domain and are likely transporting and/or interacting with the same cargo within the cell. Unfortunately, none of these proteins have three-dimensional structures deposited in the Protein Data Bank (PDB). The closest available proteins with experimentally resolved structures are: ATP13A1 (P5A subfamily), the Na^+^/K^+^-transporting ATPase (P2C subfamily), the endoplasmic Reticulum Ca^2+^-ATPase (SERCA) (P2A subfamily) and the H^+^/K^+^-ATPase (P2C subfamily). We selected the endoplasmic Reticulum Ca^2+^-ATPase (SERCA)^29^ and the very recently resolved structure of ATP13A1^33^, for modelling the active site (Fig. 2B and 2C). The SERCA ATPase has a complete conservation with ATP13A2 in the active site region of interest (Fig. S1) and was already available with several ATP analogues bound, including water molecules crystallographically resolved^29^. Our final active site models match both experimental structures very accurately (Fig. 2B and 2C for SERCA and ATP13A1, respectively).

We also note that any of the ATPases listed in Fig. 2A would be a suitable choice for modelling the active site of ATP13A2 as the Mg^2+^-binding residues in the catalytic P-domain are highly conserved among P-type ATPases, as well as the other amino acids found in the immediate active site.

### Molecular Dynamics (MD) Simulations

To probe the conformational dynamics of the protein, first, MD simulations were performed on the membrane-embedded model based on SERCA. The overall structure and in particular, the active site, was well conserved during the simulations (Fig. S5).

We focused on the Mg^2+^ ion coordination, which was initially octahedrally coordinated, coordinating: two water molecules, the side chain of the Asp513 residue (monodentate), the side chain of Asp878 (monodentate), the main chain of Thr515 (monodentate) and the γ-phosphate of the ATP. During the simulations the coordination of the Mg^2+^ ion has remained octahedral, and importantly, the binding mode of the Asp513 residue has remained monodentate. The γ-phosphate of the ATP phosphate chain was also in the correct orientation for the phosphoryl transfer to occur.

Importantly, the ATP molecule no longer preserved its original “zig-zag” conformation but immediately adopted a “straight” conformation, from the first nanosecond of the simulation (Fig. S6A). This has also been observed before in the short MD simulations of SERCA^35^. Komuro Y., *et. al* suggested that the “straight” ATP conformation is incorrect and reparameterization of the ATP molecule is needed, to ensure that the original conformation from the crystal structure ACP is preserved. We also noticed that there was a charge imbalance in the active site that could possibly cause this conformational change. The initial simulation based on the crystal structure involved a single Mg^2+^ cation only in the active site, however, we observed that a second cation constantly occupied a position very close to the α and β-phosphates (Fig. S6B). We conducted subsequent MD simulations where the ATP molecule was fixed at its original conformation. The rest of the ions, water and the protein were free to move without any constrains. In this simulation, we also observed that K^+^ ions approached the active site for the full simulation time as before, and were located in the same position where a second Mg^2+^ cation was found in the crystal structures of the homologous SERCA active site TS-like crystal structure^32^ (Fig. 3A). Throughout the duration of the MD simulations, the K^+^ ions remained a constant presence at the active site (Fig. 3B).

**Figure 3.**
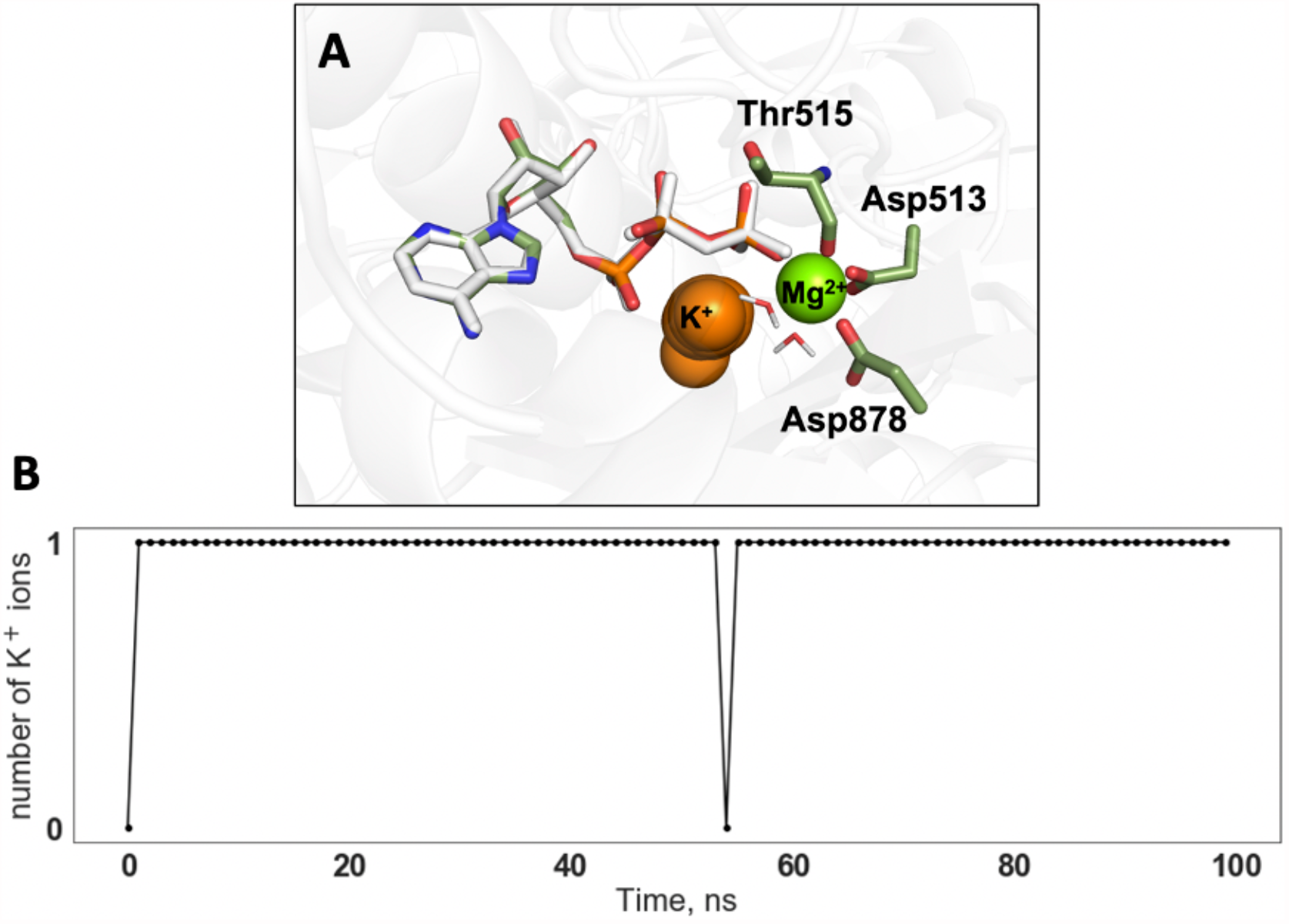
**(A)** Region of the active site where the K^+^ ion clustering is observed during the simulation (orange spheres); **(B)** Number of K^+^ ions in the active site during the duration of the 100 ns simulation.

As crystallographic structures often lack catalytically essential Mg^2+^ ions, we propose that the second Mg^2+^ ion could be needed to stabilize the catalytically active conformation of the ATP. Two Mg^2+^ ions have been resolved in the structures of homologous enzymes, however, only in TS-like states, most likely because the chain of the synthetic derivative analogue has a different charge than the catalytically active ATP.

### QM cluster Calculations

To probe the catalytically active conformation of the ATP molecule at the active site, and, specifically, the second Mg^2+^ ion coordination, we performed QM cluster calculations. We generated four different ATP-Mg^2+^ starting geometries, G1-G4 (Fig. S3), by using information from the crystal structures containing two active site cations and ADP^30,32,34^ (G1) and from preliminary results of the initial MD simulations with one Mg^2+^ in the active site (G3) or by manual initial guess (G2 and G4). All four starting geometries G1-G4 were optimized and upon convergence yielded three distinct conformations. The optimized structures of G1 and G2 (Fig. 4, green sticks) are most energetically favourable and agree particularly well with the conformation and coordination mode observed in the crystal structures containing two Mg^2+^ ions and ADP^32^ (Fig. 4, grey sticks), where the second Mg^2+^ ion is coordinating only two oxygen atoms coming from the α and β phosphate of the ATP phosphate chain, and four water molecules. Importantly, the phosphate chain in both G1 and G2 is not in a straight conformation such as in geometries G3 and G4. Conformations G3 and G4 in which the phosphate chain is “straight” (Fig. S3C and S3D) have considerably higher energy and were therefore not used in any further QM/MM calculations. Additionally, upon aligning the optimized geometries G3 and G4 with “straight” chain to the crystal structure^55^, larger deviations are clearly visible (Fig. S7C and S7D). Conformations in which the second Mg^2+^ cation is coordinated by three oxygen atoms are not favourable either (Fig. S3B) and converge to the coordination mode observed in G1 and G2, Fig. 4. This ATP conformation and coordination mode were subsequently used for QM/MM calculations to determine the corresponding reaction profile.

**Figure 4.**
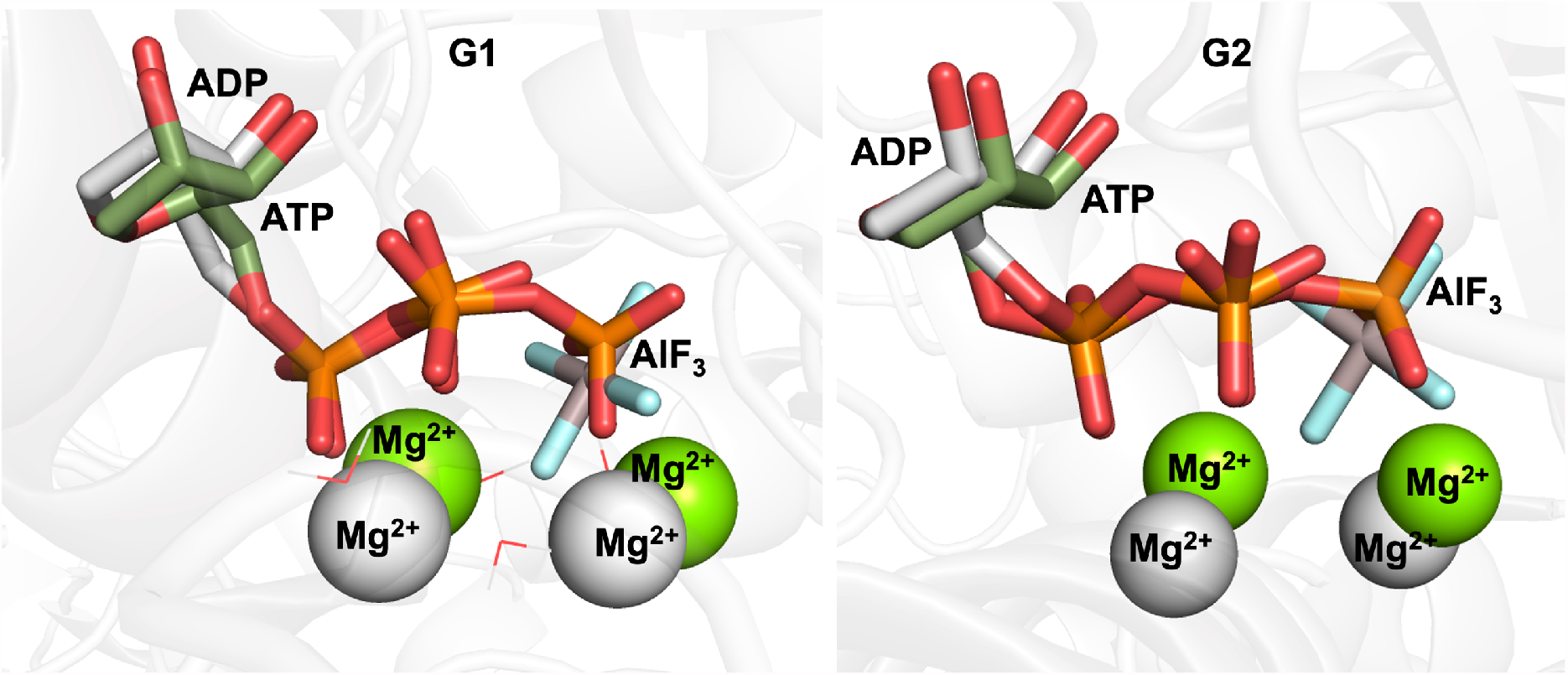
Optimized active site geometries for G1 and G2 (green sticks), aligned to the existing homologous active site crystal structure of the Na^+^/K^+^ ATPase (grey sticks, pdb: 3wgu)^32^ which is resolved with two Mg^2+^ ions (grey spheres) and an ADP molecule (grey sticks). The green spheres represent the two Mg^2+^ ions in the optimized geometries and the grey spheres represent the two Mg^2+^ ions in the crystal structure. Both structures represent the same binding mode to the second Mg^2+^ via the two oxygen atoms of the phosphate chain of the ATP.

### QM/MM Calculations

For the QM/MM calculations, six systems (P1-P6) were created using the same number and species of atoms in the MM region. However, we varied the number of atoms in the QM region to explore various effects related to the role of the number of Mg^2+^ ions, and the Lys859 and Arg686 residues for the reaction. System P1 included one Mg^2+^ ion, six water molecules, the phosphate chain of ATP, the side chain of Asp513, the side chain of Asp878, Thr515, the side chain of Lys859, Arg686, Gly516 and the main chain of Lys514 in the QM region. System P2 contained the same number and atom species but also a second Mg^2+^. P3 and P4 (Table S1) contain one Mg^2+^ ion in the active site and Lys859 or Arg686, respectively, are removed from the QM region and their MM atomic charges are set to 0. P5 and P6 contain two Mg^2+^ ions in the active site (Table S1) with Lys859 or Arg686, respectively, electrostatically removed from the system as detailed above. Reaction scans were performed on all six systems, P1-P6, to determine the potential energy barriers. We performed forward and backward scans until we observed consistent energy profiles (Fig. 5A) and the energy minimum was obtained. Both scans show a good convergence. For P1, in the presence of one Mg^2+^ only, the barrier height is ∼12.5 kcal mol^-1^. During the phosphoryl transfer reaction, the ion is octahedrally coordinated via two water molecules, the γ-phosphate of the ATP phosphate chain, the carboxylate side chains of Asp878 and Asp513 (in a monodentate mode) and the main chain carbonyl of Thr515 (Fig. 5B). The barrier height for P2, containing two Mg^2+^ ions was calculated to be ∼7.0 kcal mol^-1^. The second Mg^2+^ ion coordinates four water molecules and two oxygen atoms coming from the α and β phosphates of the ATP chain. Importantly, it does not coordinate a third oxygen atom from the phosphate chain.

**Figure 5.**
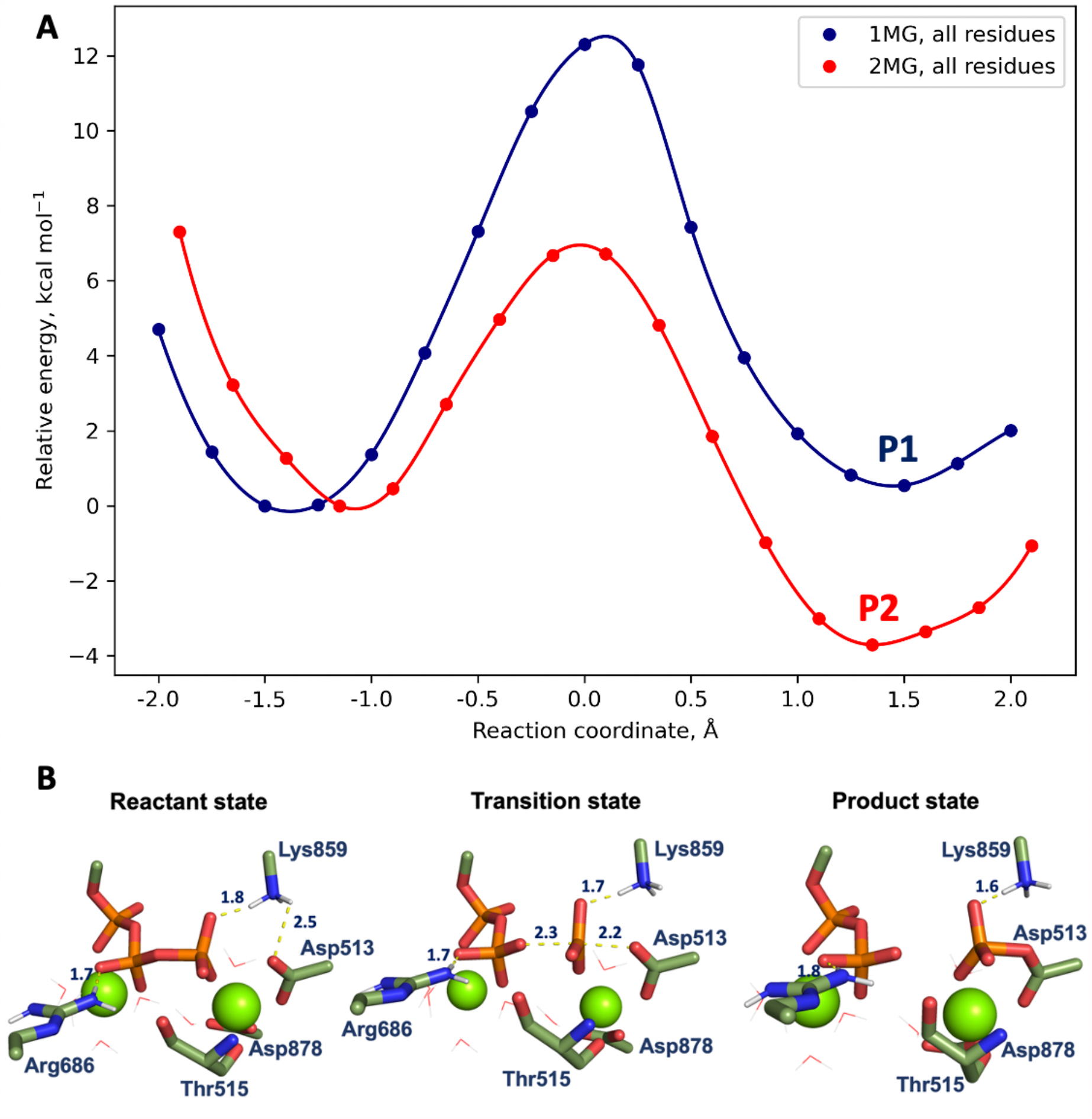
**(A)** Reaction coordinate scan with 1 (blue) and 2 (red) Mg^2+^ ions in the active site of ATP13A2. The reactant state is set to 0. **(B)** Reactant, transition state and product state during the ATP cleavage reaction with 2 Mg^2+^ ions in the active site are shown (green sticks). Distances are shown between the Lys859 and Arg686 and the ATP molecule. O_2D_–P_G_–O_3B_ distance is shown in the TS state. Hydrogens on the rest of the amino acids are not shown for clarity. Throughout the RCS, the Mg^2+^ ion coordinating Asp513 is always bound to two water molecules and the Mg^2+^ ion which is coordinating the α and β-phosphates of the ATP chain, coordinates four water molecules.

Kinetic studies of the Ca^2+^-transporting Sarcoplasmic Reticulum (SR), which has a fully conserved active site with ATP13A2, report a rate constant of 225 s^-1^ for the phosphorylation of the wild type enzyme by ATP in equilibrium conditions^58^. This experimental report agrees with the work of Petithory *et. al*^59^, which reports a rate constant for formation of the phosphorylated enzyme of 220 s^-1^. This corresponds to a barrier height of ∼14.3 kcal mol^-1^ using the Eyring equation at 298K. Inesi *et. al* reported experiments with a somewhat slower rate (100-150 s^-1^)^60^. Our barrier height obtained for the one Mg^2+^ case, is in excellent agreement with these kinetic experiments. All experimental work agrees that the enzyme phosphorylation reaction is fast, and it is not the rate limiting step, with the expected barrier ranging from ∼14-15 kcal/mol^-1^.

We find that the phosphoryl transfer reaction proceeds as a one-step reaction, without any stable pentavalent phosphate intermediate. The reaction pathway in phosphoryl transfer reactions can be classified as associative, dissociative or concerted^61,62^. In the associative pathway, the attacking nucleophile approaches the phosphorus atom, decreasing the bond length between the attacking nucleophile and the reactive phosphorus while the bond to the leaving group simultaneously increases.

This is generally a lower energy pathway for biological phosphate reactions. Alternatively, in the dissociative pathway, the leaving group departure precedes the nucleophilic attack. In the concerted pathway, partial bond formation and bond breaking occur simultaneously. Based on the evolution of the distances between the attacking nucleophile and the reacting phosphate, and the phosphate and the leaving group, the reaction pathway can be classified. In this ATP cleavage mechanism of ATP13A2, there is an energetically stable leaving group, a diphosphate anion, and a dissociative pathway could be expected to potentially be more favourable^60^. To confirm, we considered the distances between O_2D_-P_G_ and P_G_-O_3B_ (Fig. 6A). Our results show that the path from the reactant state (RS) to the product state (PS) follows an associative pathway in the protein environment, for both one and two Mg^2+^ cases. Furthermore, we note that while the reaction can follow an associative pathway, the TS itself does not have to be associative^62,63^. We calculated the bond order in the transition state and obtained Wiberg bond indices for the P_G_-O_2D_ and P_G_-O_3B_ bonds of 0.162 and 0.125. We therefore found that while the reaction clearly follows an associative pathway (Fig. 6A), the TS itself corresponds to a loose, dissociative structure, where both P-O bonds are already broken (Fig. 6B). This is consistent both with computational work demonstrating an associative path^61^ and with experimental findings that can capture the loose TS.^64^

**Figure 6.**
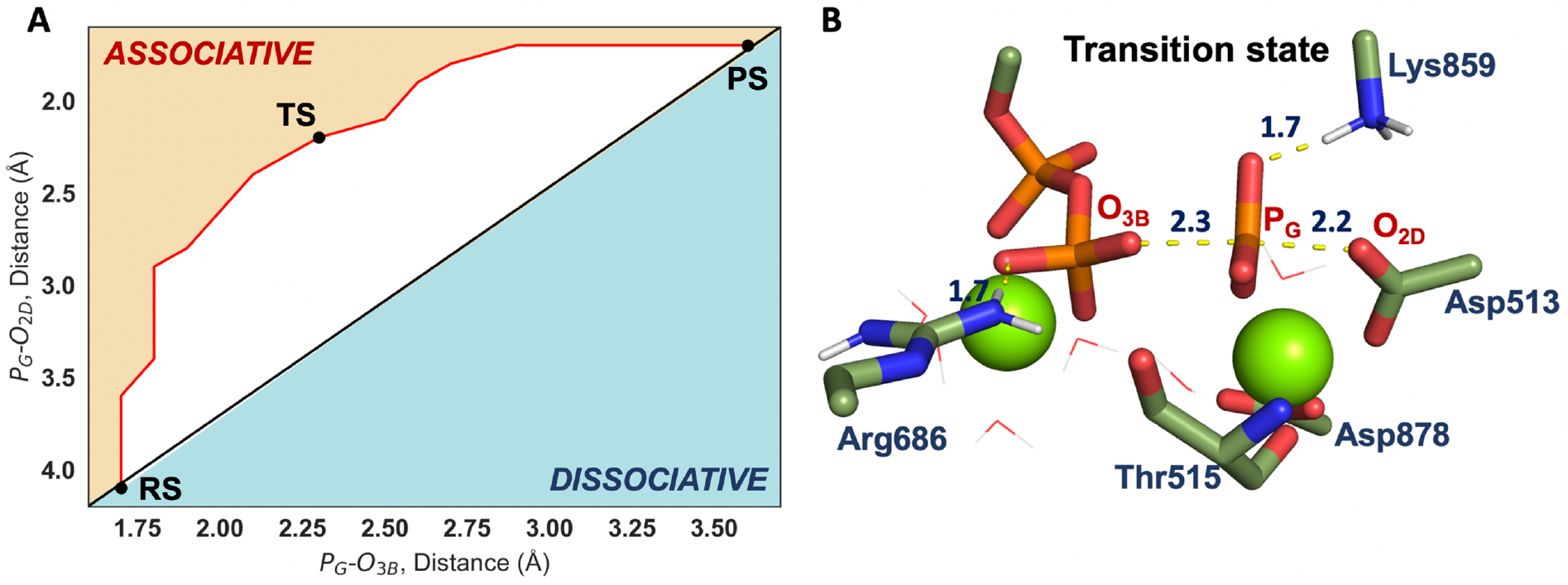
**(A)** The path adopted for the ATP cleavage is associative, based on the distance evolution from the reacting nucleophile O_2D_ to P_G_ and P_G_-O_3B_ (leaving group), respectively. **(B)** Details of the transition state structure for the reaction. Key residues at the active site (sticks and spheres for Mg^2+^) and relevant distances (yellow dashed lines) are shown.

To quantify the effects of the interactions between the ATP and Lys859 or Arg686 (P3-P6), we obtained converged Reaction Coordinate Scans by eliminating these interactions (Fig. 7). This allows us to observe the reaction coordinate without the stabilizing electrostatic interactions formed between Arg686 and the β-phosphate of the ATP chain; and between Lys859 and both the γ-phosphate and the Asp513. Without Arg686, the barrier height increases by more than 5.0 kcal mol^-1^ (Fig. 7), which is very significant in terms of time scales. Without Lys859, which forms key electrostatic interactions to the side chain of Asp513 and the γ-phosphate of ATP (Fig. 5B), the reactant state is considerably higher in energy than in the cases when Lys859 is present in the system. While this results in a lower apparent barrier, the reactant state is very destabilized and the ATP binding is likely impaired, which might have additional consequences for the enzyme function and stability. Our observation that the Lys859 residue is very important for the phosphoryl transfer reaction is supported by the kinetic analysis of mutants of the homologous Ca^2+^-ATPase Sarcoplasmic Reticulum by Sorensen *et. al* ^58^. This work shows that the rate of ATP binding and subsequent phosphoryl transfer in the Lys684Arg mutant (corresponding to Lys859Arg in ATP13A2), was reduced 50-fold, relative to wild type, thus indicating the importance of this residue. This information combined with our analysis shows that the structural effects are also very important in case of the Lys859Arg substitution, as the electrostatic factors here are unchanged, which we predicted to lower the barrier, therefore the overall geometrical changes significantly slow down the reaction. The transition states for system P5 and P6 are displayed in Fig. S8.

**Figure 7.**
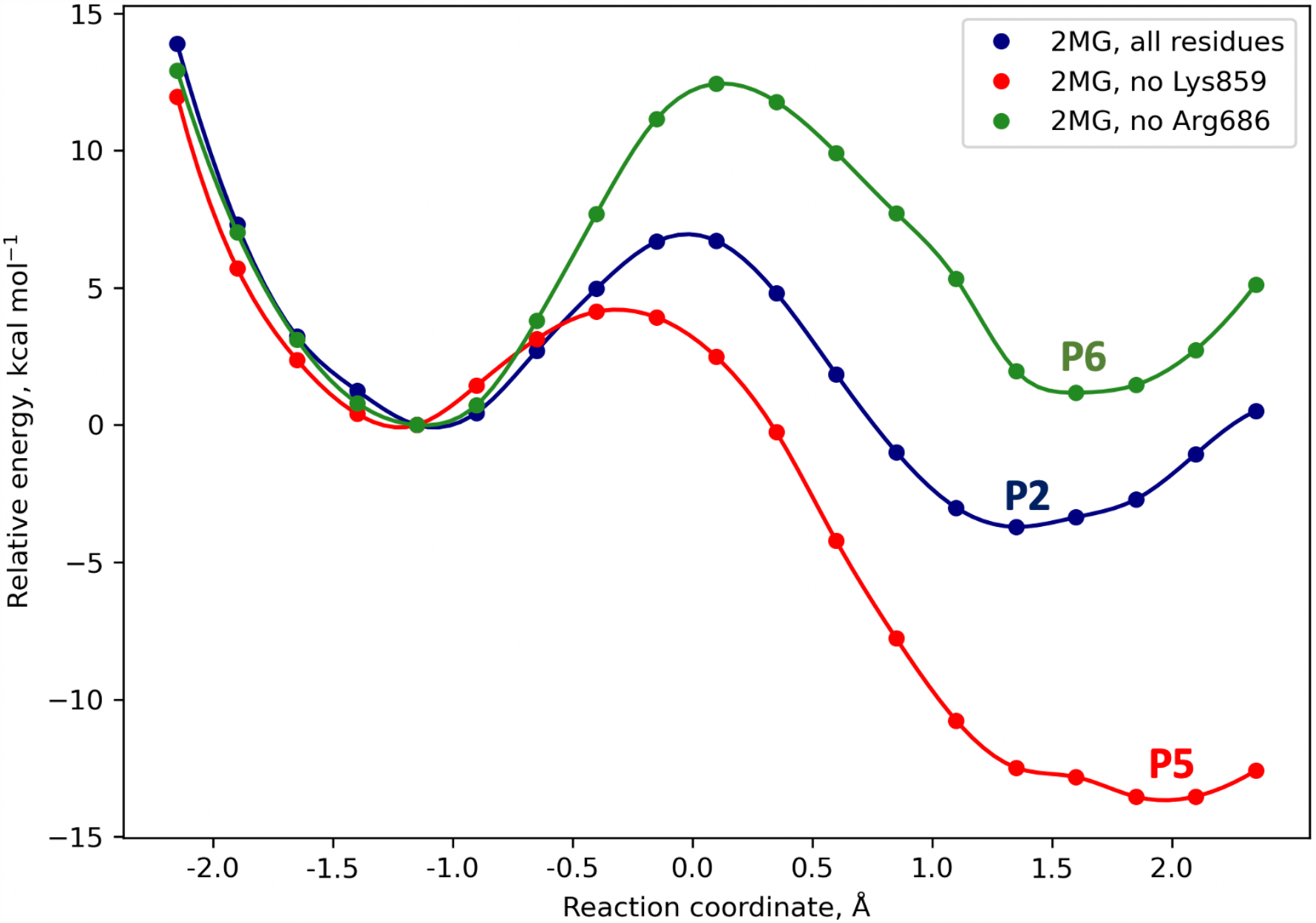
Reaction coordinate scans for the active site of ATP13A2 containing two Mg^2+^ ions. The RCS depicted in red represents the phosphate transfer reaction without Lys859 (P5). The RCS in green represents the phosphate transfer reaction without Arg686 (P6). The blue RCS shows the reaction in the active site when all amino acids required for the normal enzymatic activity, are present.

### Binding Pockets and Transmembrane binding Analysis

While the P5A ATPase ATP13A1 has been shown to be a Mn-transporter^65^, it has recently been demonstrated that ATP13A2 is strongly implicated in polyamine export^23^. It has been known that P5B ATPases such as ATP13A2 likely transport different cargo from ATP13A1 due to major differences in the transmembrane domain sequence conservation^66^. To identify binding regions on the surface of ATP13A2, pocket analysis was performed on twenty equidistantly spaced frames from the MD trajectories of the modelled protein. The four biggest pockets were analysed for each frame. Not surprisingly, for most of the frames the biggest pocket in terms of surface area is the one where ATP binds in the N-domain (Fig. 8A). It is expected that this pocket will be conserved for all homologous P-type ATPases due to the highly conserved ATP binding region and mechanism in the active site.

**Figure 8.**
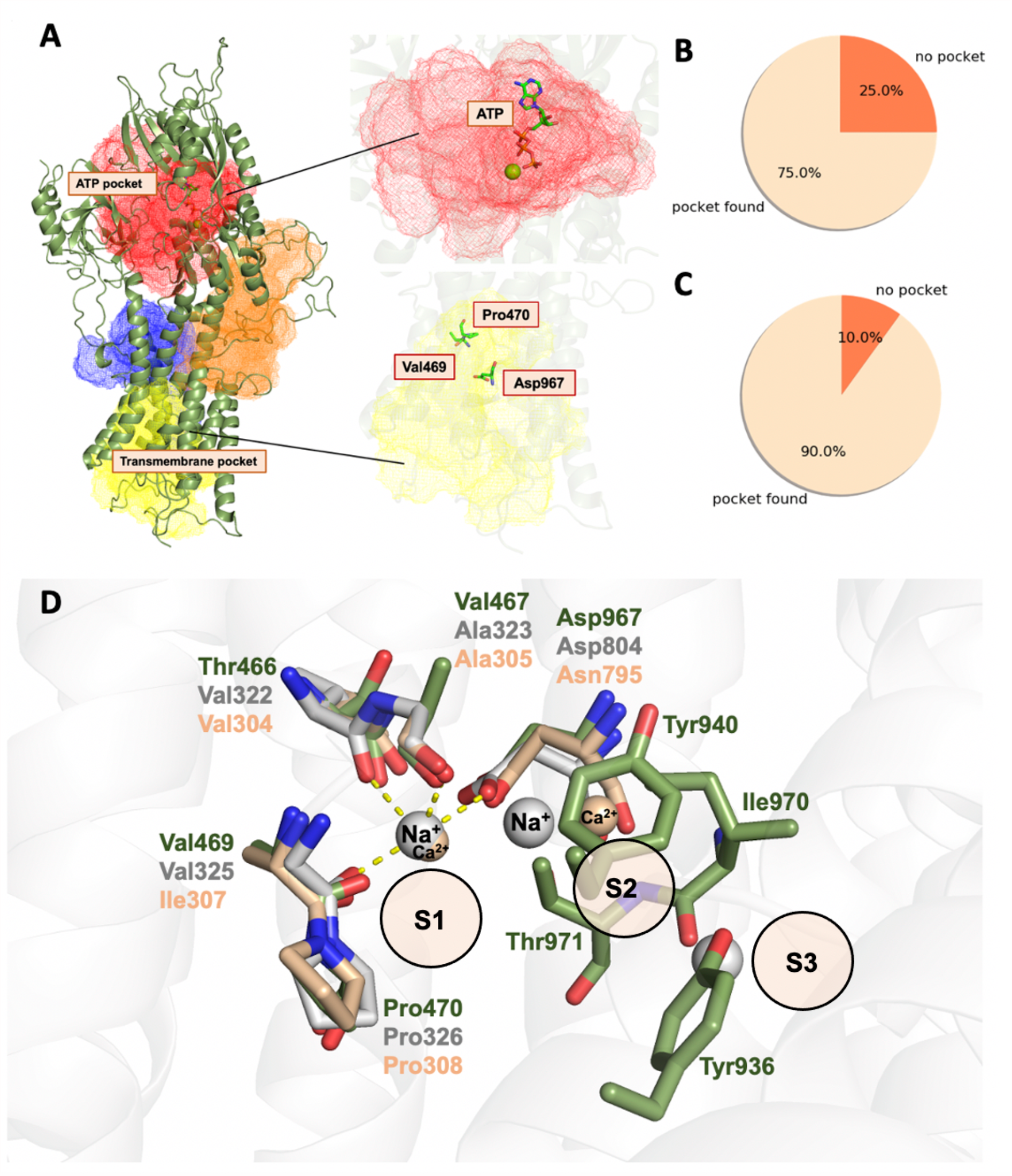
**(A)** The two main binding pockets (red mesh and yellow mesh) found consistently on the surface of the homology model of ATP13A2 (green cartoon). **(B)** Frequency of ATP-pocket occurrence calculated from the 20 frames of the unconstrained MD simulations. **(C)** Frequency of transmembrane pocket occurrence calculated for the same frames. **(D)** Sequence conservation in the inorganic ion-binding region in the transmembrane between the Na^+^/K^+^-ATPase (grey sticks), SERCA (wheat sticks) and ATP13A2 (green sticks). S1, S2 and S3 stand for binding sites 1, 2 and 3.

Interestingly, the second biggest pocket, which appears consistently, was identified in the transmembrane region (Fig. 8A), more specifically, where inorganic ion binding has been observed in the crystal structures of homologous enzymes such as the P2C Na^+^/K^+^-ATPase^31^ and the P2A ATPase SERCA^29^. We call this spatial region Site 1 (S1), Fig. 8D. For the Na^+^/K^+^-ATPase, the specific amino acid scaffold surrounding the three Na^+^ ions which bind in the transmembrane domain consists of: Val322, Ala323, Val325, Pro326, Glu327, Tyr771, Thr774, Ser755, Asn776, Glu 779, Asp804 and Gln924. From this sequence motif we observe conservation in ATP13A2 for the amino acids that bind one of the inorganic ions in S1 (Fig. 8D). In ATP13A2, these correspond to: Val469, Pro470 and Asp967 (Fig. 8D). The remaining amino acids that coordinate the additional two Na^+^ ions in the Na^+^/K^+^-ATPase (Site 2 and 3, Fig. 8D) are not conserved in ATP13A2. Val469 and Pro470 in S1 are also conserved between ATP13A2 and SERCA, which transports Ca^2+^ (Fig. 8D), but the amino acids coordinating the second ion in SERCA and Na^+^/K^+^-ATPase (S2 and S3) are not conserved in ATP13A2. Considering the amino acid conservation in the ion-binding region of the transmembrane (Fig. 8D), and overall similar shape of the ion-binding scaffold (Fig. 8D) it is possible that ATP13A2 also can bind and transport one inorganic ion, although this is unlikely to be either Na^+^ or Ca^2+^. Additionally, this transmembrane pocket was consistently (90% of the time of the analysed frames) found within 1.5 Å of ion-binding amino acids (Val469, Pro470, Asp967) in the S1 region, which additionally supports the idea that the protein binds cargo in this area. It is important to note that the surface area of this pocket is considerably larger than expected from an ion binding site alone – the average size of the transmembrane pocket is 2264.61 Å^3^. This suggests that this region of the protein additionally could interact with a much bigger substrate, likely in S2 and S3 region which is not conserved between the Na^+^/K^+^-ATPase or SERCA, Fig. 8D.

Two additional pockets also appeared frequently, however, not as consistently as the ATP pocket and the main transmembrane pocket. These include a considerably smaller pocket in the upper part of the transmembrane (Fig. S4) and an additional pocket around residues 985-995 and 800-807 (Fig. S4).

## Conclusions

In this work we present a structural model of the P5B enzyme ATP13A2 which has been complemented with MD, QM cluster and QM/MM calculations. This has allowed us to find an accurate conformation for the catalytically competent ATP structure and a reliable position for the second Mg^2+^ ion with respect to the ATP and the other Mg^2+^ active site cation. Using this information, we have subsequently calculated the barrier height for the phosphoryl transfer with one and two Mg^2+^ ions. Additionally, we present the first quantitative analysis of the role of Arg686 and Lys859 on the barrier height of the ATP cleavage and phosphoryl transfer. This work can suggest how missense mutations close to important active site interactions in the respective catalytic domains, can have diminishing effect on the catalytic activity of the enzyme.

Additionally, we have analysed the surface of the protein for binding pockets and found two pockets which occur consistently. The pocket which appeared most consistently was found in the transmembrane domain of the protein. From the sequence analysis performed and the binding pocket calculations, we suggest that ATP13A2 also likely binds a substrate in this part of the transmembrane, other than an inorganic ion, which is consistent with the big size of the calculated pocket in this part of the transmembrane.

## Acknowledgment

We are grateful to Tamás Földes for numerous discussions and also for his kind advice and help to create better visualisations for this work. We would also like to thank Attila Csikász-Nagy for initial discussions. ER acknowledges funding from the ERC (project 757850 BioNet). TM acknowledges funding from the Agency for Science, Technology and Research (A*STAR) Singapore Research Attachment Programme (ARAP) and King’s College London’s Centre for Doctoral studies. We acknowledge the use of the research computing facility at King’s College London, Rosalind (https://rosalind.kcl.ac.uk).

## ASSOCIATED CONTENT

Supporting Information including

- Figures S1-8, Alignments, data from the MD, QM structures before and after optimization, Transition states for systems P5 and P6 and additional pocket details.
- Tables S1-3, Energy of the optimized geometries, details of QM/MM systems and additional pockets information from the Binding Pocket calculations.

## Author Information

### Present Addresses

Department of Chemistry, Faculty of Natural & Mathematical Sciences, King’s College London, 7 Trinity St, London SE1 1DB, United Kingdom

Department of Materials Science and Chemistry, Institute of High Performance Computing, Agency for Science, Technology and Research (A*STAR), 1 Fusionopolis Way, 16-16 Connexis (North Tower), Singapore 138632

Department of Physics and Astronomy, Faculty of Maths & Physical Sciences, University College London, Gower Street, London WC1E 6BT, United Kingdom

### Notes

The authors declare no competing financial interest.

## TOC Graphic

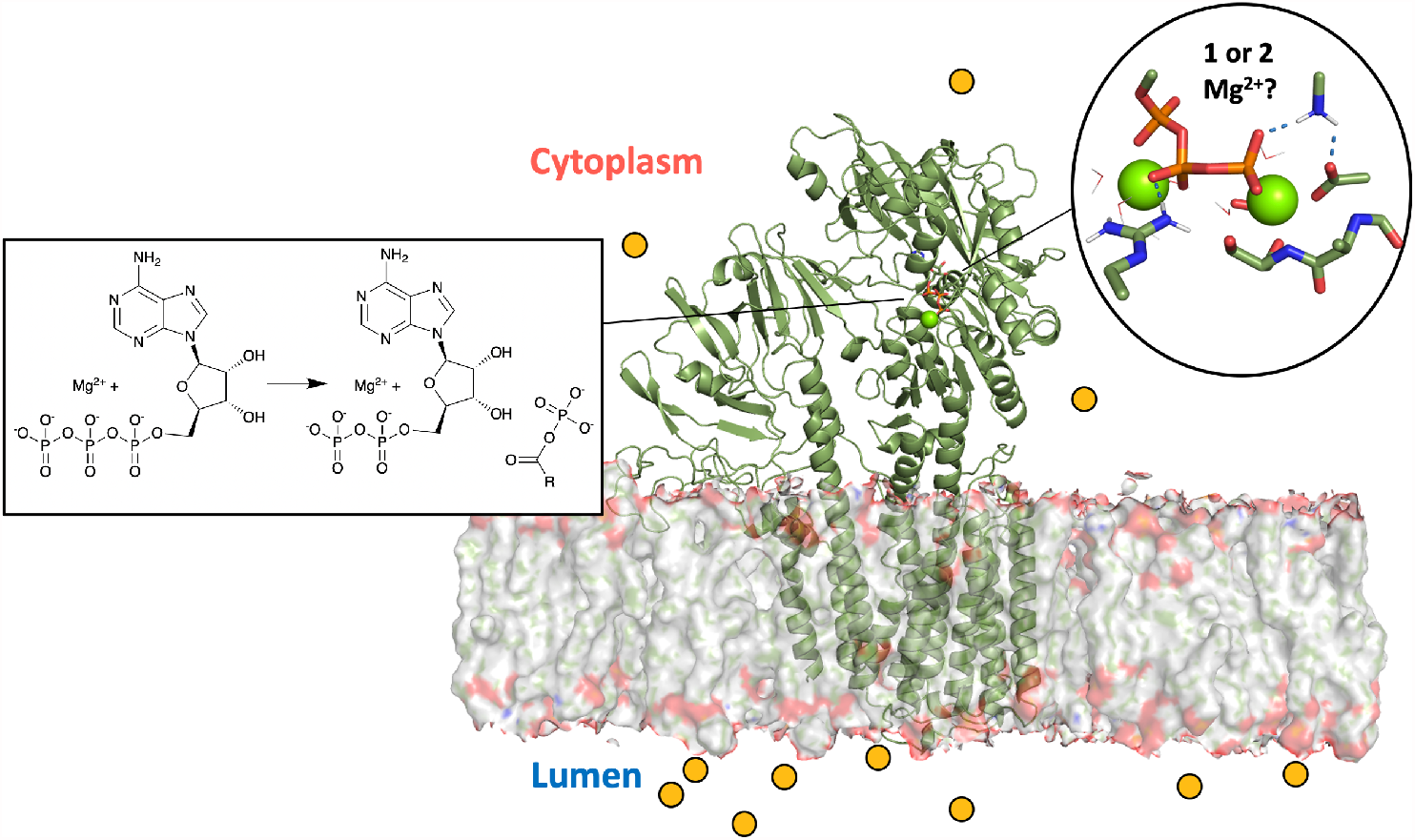

