## Supplementary Information for "Structural Dynamics and Catalytic Mechanism of ATP13A2 (PARK9) from Simulations"

### Homology Modelling

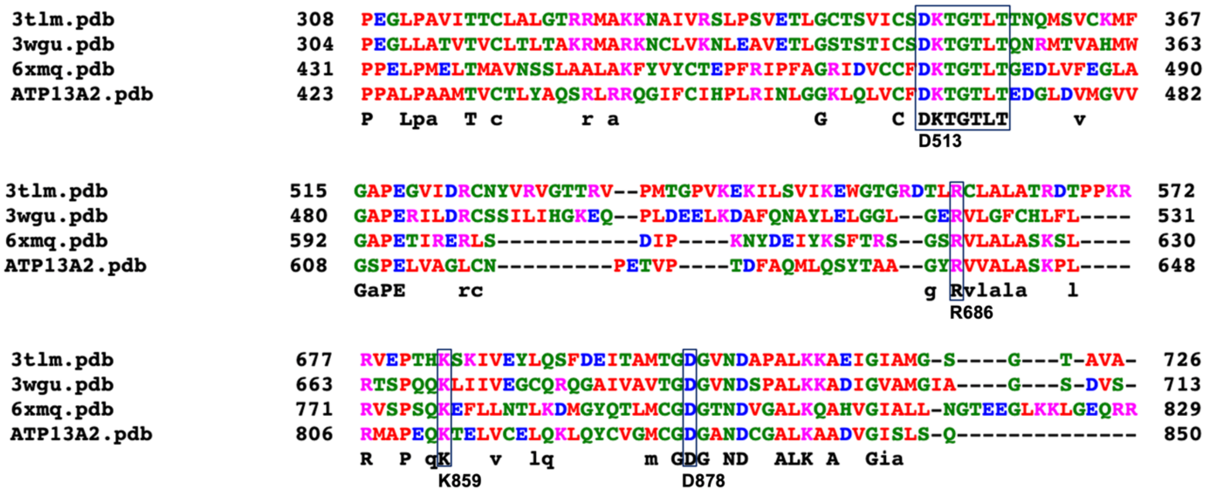


**Figure S1.** Alignment of the sequences of the most homologous proteins to ATP13A2 which have crystallographic structures deposited in the Protein Data Bank (PDB). Only regions of interest in this work are shown. The **DKTGTLT** motif is conserved among all protein structures used in this work. The $Mg^{2+}$-coordinating active site residues D513 and T515, correspond to **D**K**T**GTLT in this motif. The active site residues R686 and K859 are also strictly conserved among all P-type ATPases used in this work. D878 is the second aspartate amino acid which coordinates one of the $Mg^{2+}$ ions during the hydrolysis of ATP. The proteins aligned here are: SERCA (PDB code: 3tlm)[1], $Na^+/K^+$- ATPase (PDB code: 3wgu)[2], ATP13A1 (PDB code: 6xmq)[3] and the homology model of ATP13A2. The numbering below the conserved residues corresponds to the number of the residue in the sequence of ATP13A2 for Homo Sapiens (Uniprot[4] code: Q9NQ11).

**QM Cluster Calculations**

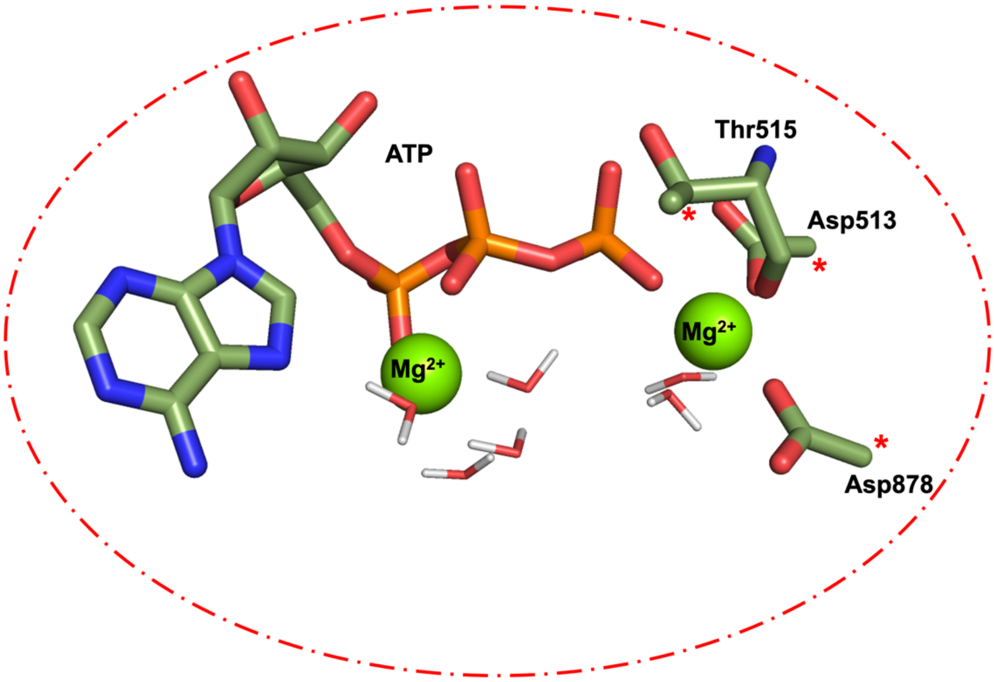


**Figure S2.** All atoms included in the QM cluster geometry optimizations. Red asterisk marks where an amino acid residue has been truncated, as well as which atom was frozen during the geometry optimization. Each calculation contained six water molecules, two $Mg^{2+}$ cations, the ATP moiety and the three amino acids depicted.

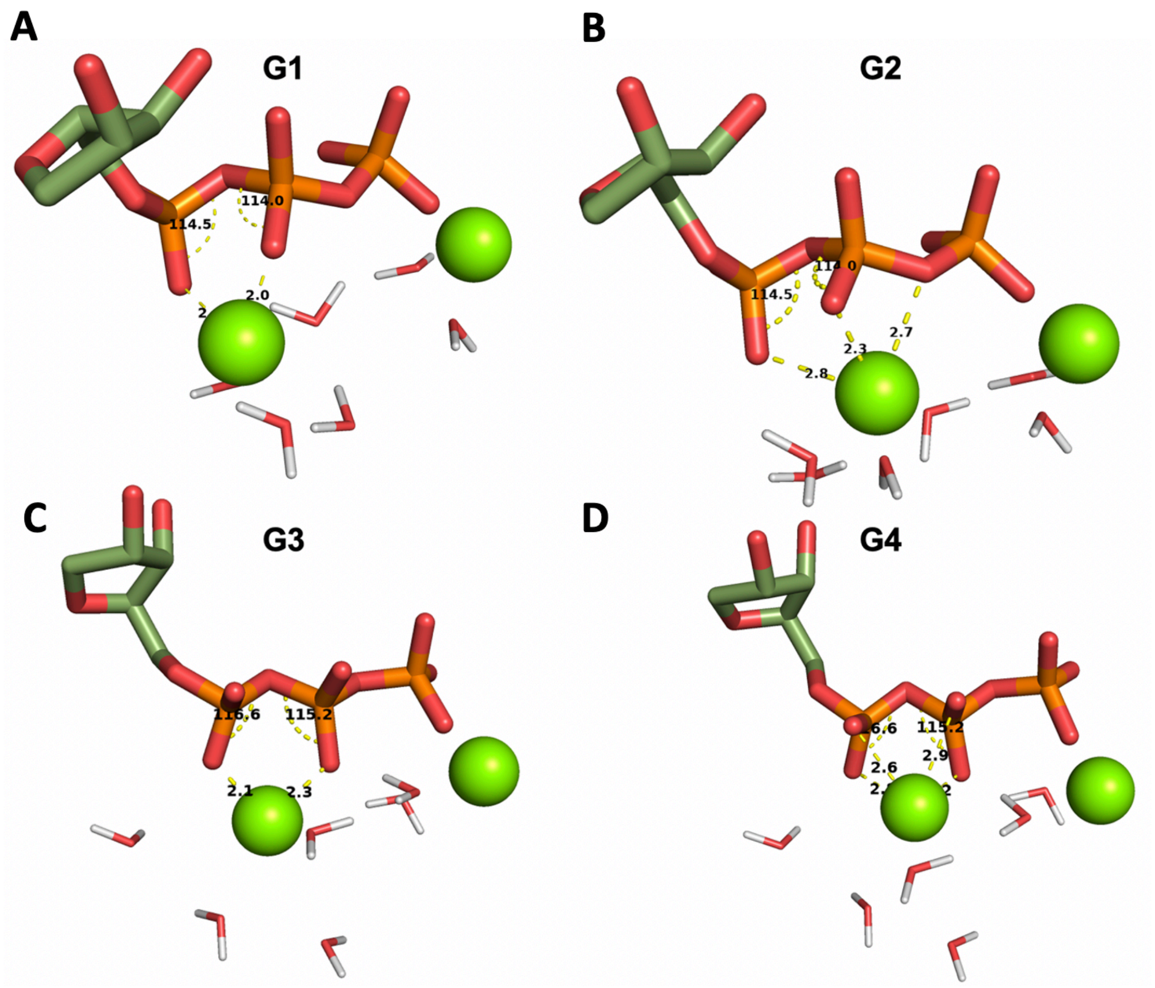


**Figure S3.** Starting geometries G1-G4 for the QM cluster calculations (before optimization). Starting structure G1 **(A)** was created based on alignment with the crystal structure resolved with ADP and 2 $Mg^{2+}$ ions (PDB code: 3wgu)[2]. Structure G2 **(B)** exhibits an alternative binding but has the same conformation as **(A)**, here the second $Mg^{2+}$ ion is coordinating three oxygen atoms instead of two. G3 **(C)** was taken from the last frame of the original MD simulation (after 100 ns) and G4 **(D)** represents a more unorthodox geometry that has not been observed by us in any P-type ATPase crystal structures but was generated to explore different possibilities. The “straight” conformation of the phosphate chain G3 and G4 in **(C)** and **(D)** is unfavorable energetically.

**QM/MM Reaction Coordinate Scans (RCSs)**

Six systems were created (P1-P6) which were initially minimized for 1000 steps. The minimized structure of each system was supplied as a starting point for the Reaction Coordinate Scan (RCS). P1 refers to the full system containing all amino acids described to be contained in the QM region, including Arg686 and Lys859, with one $Mg^{2+}$ in the active site. P2 refers to the full system containing all amino acids described to be contained in the QM region, including Arg686 and Lys859, with two $Mg^{2+}$ ions in the active site. P3 and P4 have one $Mg^{2+}$ ion in the active site and Arg686 or Lys859 missing, respectively. P5 and P6 have two $Mg^{2+}$ ions in the active site with Arg686 or Lys859 missing, respectively. The aim of obtaining a converged RCS for each system P1-P6 was to show the effect of Lys859 and Arg686, as well as the profile of the wild type system where all amino acids are present and not mutated.

**Table S1.** Six QM/MM systems (P1-P6) were initially minimized, and subsequently QM/MM reaction coordinate scans were run. The overall charge and atoms of each QM region, as well as the number of $Mg^{2+}$ ions in each active site are shown below. To test the effects of residues missing from the active site, we neutralized the atomic charges of selected residues as indicated.

| System | Number of $Mg^{2+}$ | Overall Charge | Residues Missing |
|---|---|---|---|
| **P1** | 1 | -2 | none |
| **P2** | 2 | 0 | none |
| **P3** | 1 | -3 | Lys859 |
| **P4** | 1 | -3 | Arg686 |
| **P5** | 2 | -1 | Lys859 |
| **P6** | 2 | -1 | Arg686 |

**Pocket Analysis**

Twenty equidistant frames from the 100 ns unconstrained MD simulation were extracted. The twenty biggest pockets were calculated in each frame and the four biggest pockets in terms of surface area were further analyzed. An ATP pocket was found in 75% of all frames and a transmembrane pocket, within 1.5 Å of residues Val469, Pro470 and Asp967, was found 90% of the time. The size of each pocket in the corresponding frame is listed in Table 2. Two other pockets were found consistently, and their location is shown in Figure S4. The blue pocket is relatively small in surface area and is found exclusively in the first 25 ns of the simulation, whereas the orange pocket is found in the later part of the simulation (after 25 ns).

**Table S2.** Frames extracted from the 100 ns unconstrained MD simulation and the size of two biggest pockets found.

| Frame | ATP pocket size (Å$^3$) | Transmembrane pocket size (Å$^3$) |
|---|---|---|
| **50** | 3750.0 | 1278.0 |
| **100** | **Not found** | 975.0 |
| **150** | 745.0 | **Not found** |
| **200** | 3519.0 | **Not found** |
| **250** | 2907.0 | 1541.0 |
| **300** | **Not found** | 2197.0 |
| **350** | 1093.0 | 1045.0 |
| **400** | **Not found** | 2998.0 |
| **450** | 1035.0 | 1818.0 |
| **500** | **Not found** | 1928.0 |
| **550** | 1725.0 | 2110.0 |
| **600** | 5576.0 | 3185.0 |
| **650** | 6687.0 | 1731.0 |
| **700** | 6743.0 | 4021.0 |
| **750** | 5621.0 | 3838.0 |
| **800** | 5067.0 | 3508.0 |
| **850** | 5867.0 | 2901.0 |
| **900** | 12411.0 | 1886.0 |
| **950** | 14078.0 | 1835.0 |
| **1000** | **Not found** | 1968.0 |

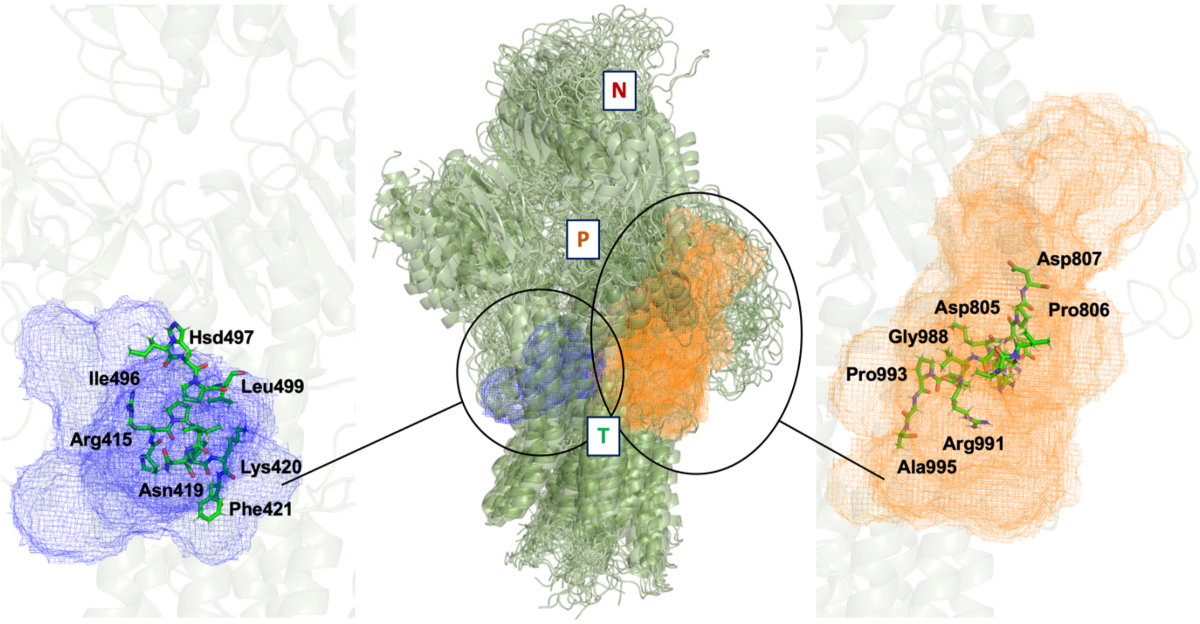


**Figure S4.** Two additional pockets (blue mesh and orange mesh) which were found on the surface of the ATP13A2 homology model (green cartoon).

### Molecular Dynamics (MD) Simulations

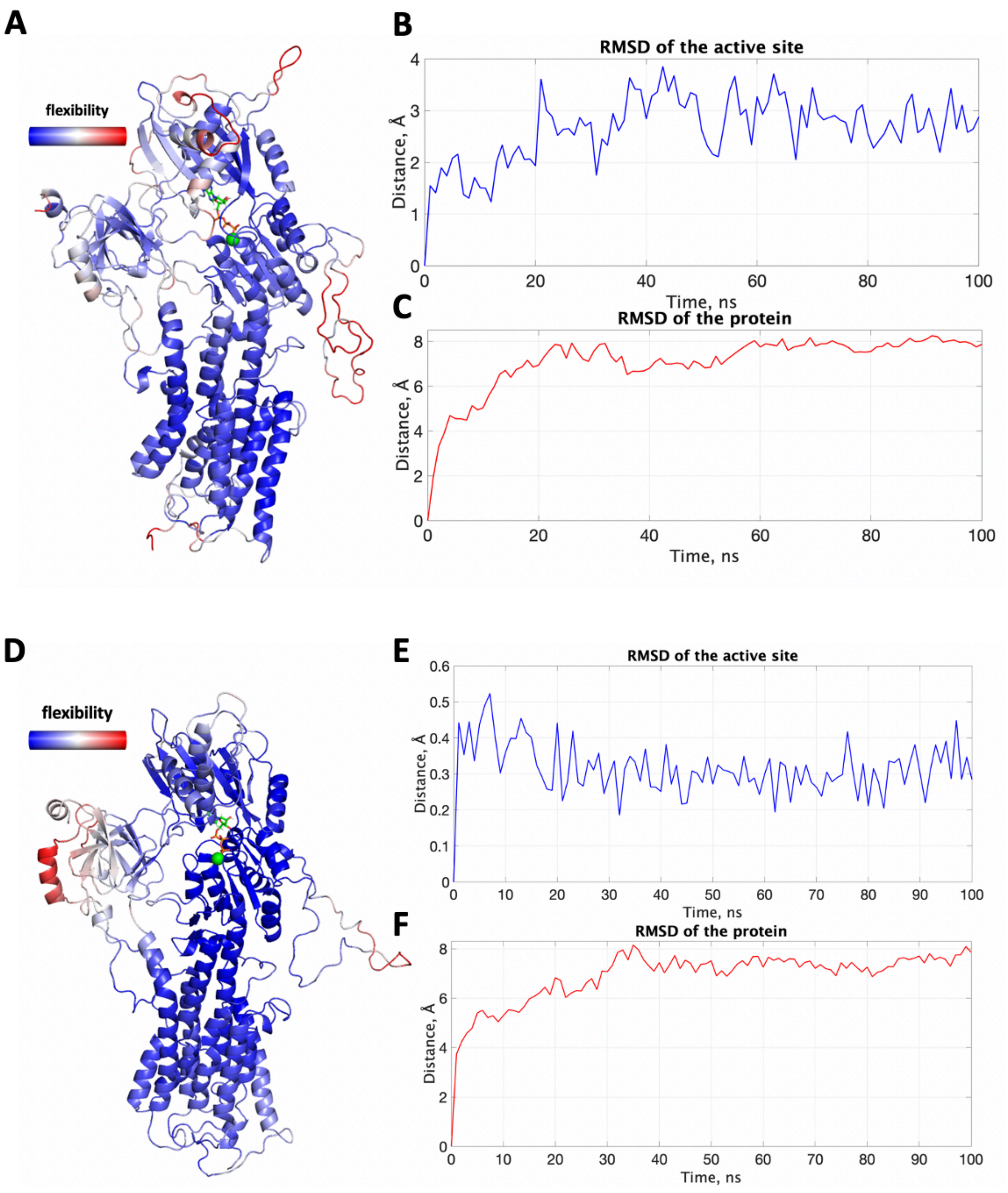


**Figure S5. (A)** The protein colored by flexibility in the unconstrained MD simulation. Red regions signify more flexible parts while blue the more rigid regions. The membrane and water molecules/ions are not included in this visualization for clarity. The RMSD of the active site features Asp513, Thr515, Asp878, the $Mg^{2+}$ cation and the ATP molecule. **(B)** shows the RMSD of the active site alone and **(C)** the RMSD of protein. **(D), (E)** and **(F)** show the same parameters for the second simulation where the ATP molecule was fixed to its original coordinates.

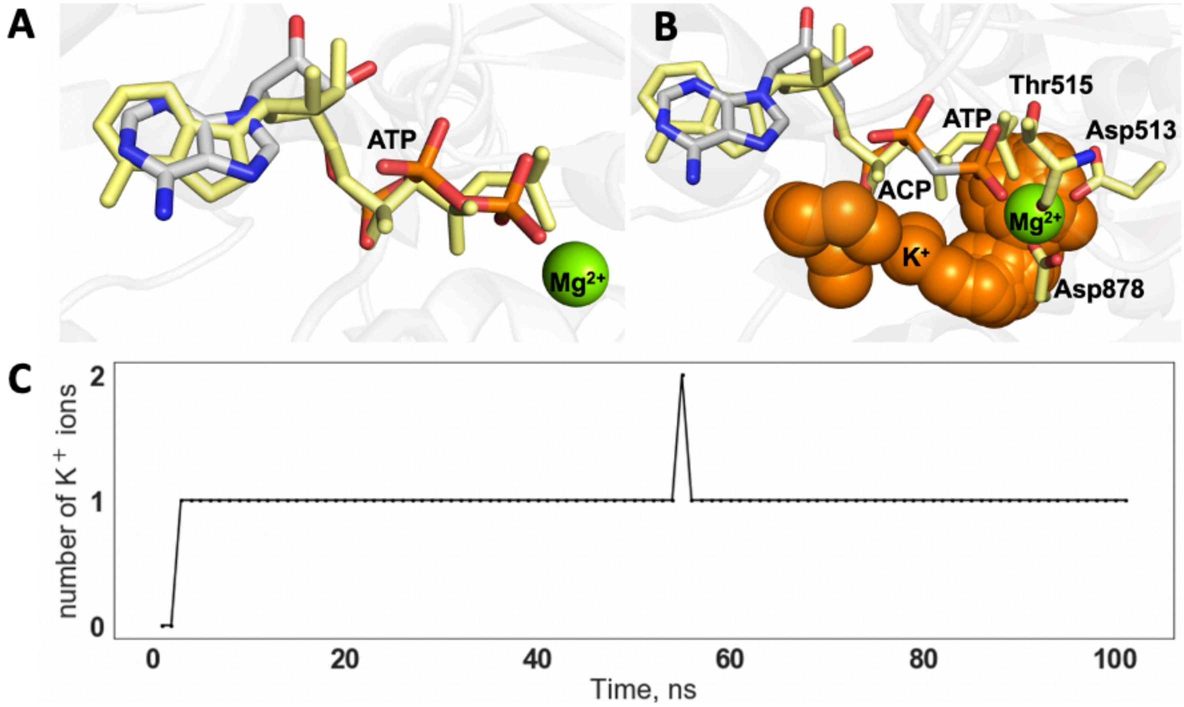


**Figure S6. (A)** Conformation of the ATP molecule (grey sticks) at the start of the MD simulation and after 1 ns (yellow sticks). The 'straight' ATP conformation (yellow sticks) adopted after the start of the simulation is a result of the parametrization of the ATP in the force field used. It does not correspond to the conformation observed in the crystal structures (the resolved molecule in crystal structures is ACP) and is energetically not favorable which was later shown in this work. **(B)** Region of the active site where the $K^+$ ion clustering is observed during the unconstrained simulation (orange spheres). The constant presence of the $K^+$ ions **(C)** suggests insufficiency of positive charge in the active site. Since the ATP has not preserved its original "zig-zag" conformation, the $K^+$ ions cluster ununiformly. **(C)** Number of $K^+$ ions in the active site throughout the 100 ns unconstrained simulation.

**QM Cluster Calculations**

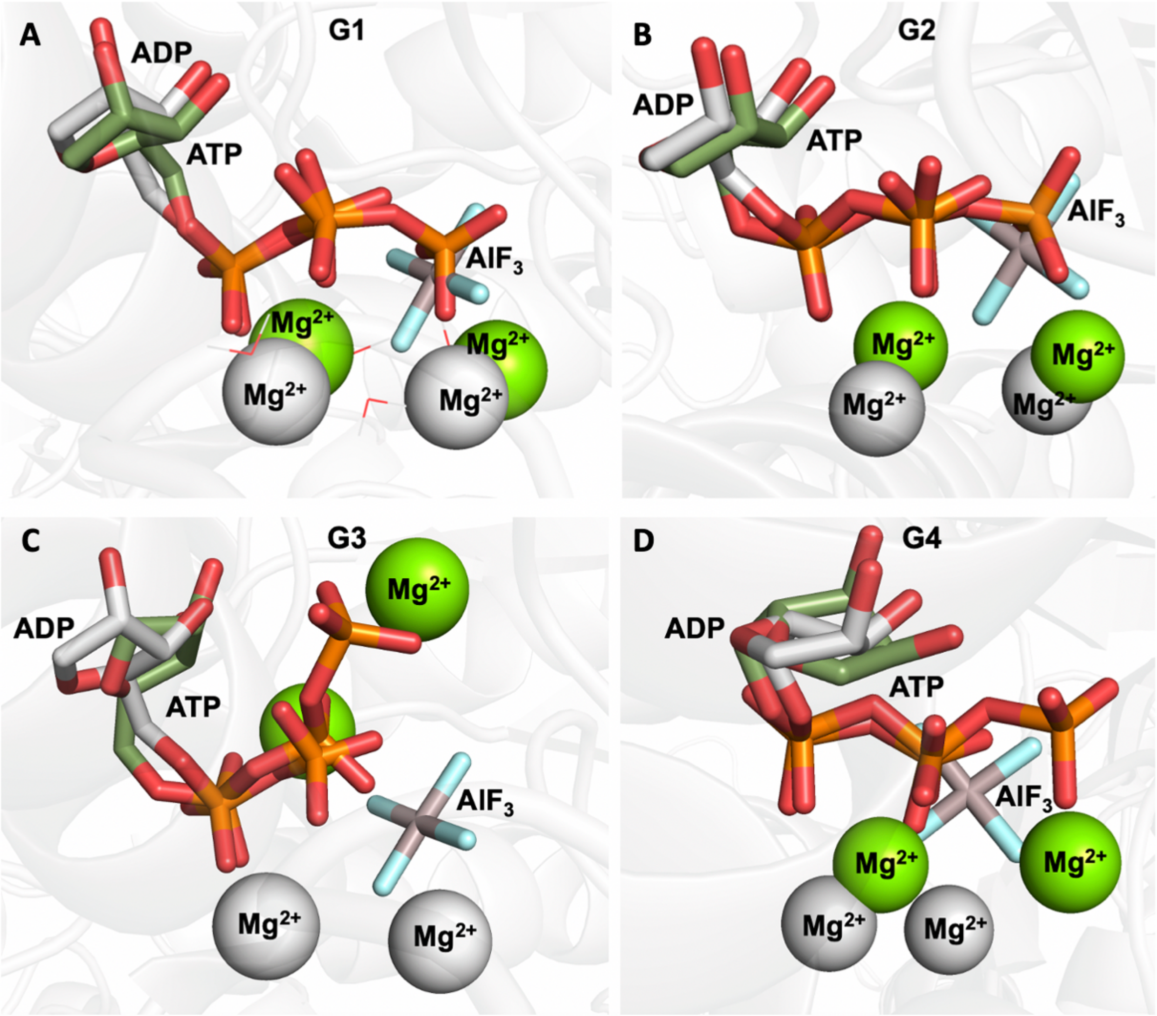


**Figure S7.** The four optimized geometries (green sticks) aligned to the crystal structure (grey sticks) resolved with ADP and two $Mg^{2+}$ ions[2]. The conformation of G1 in **(A)** and G2 in **(B)** agree very well with the crystal structure of the transition state, unlike the "straight" phosphate chain conformations of G3 and G4 **(C)** and **(D)**, respectively.

Starting Geometries G1-G4 (Fig. S3) were optimized and upon convergence yielded three distinct conformations. G1 and G2 optimize to the same type of conformation and coordination mode. All optimized geometries with straight phosphate chain (G3 and G4) are energetically unfavourable (Table S3) and were therefore not used in any further QM/MM Reaction coordinate scans (RCSs).

**Table S3.** Energy of the QM cluster optimized structures.

| Optimized conformation | Energy (kcal/mol) |
|---|---|
| G1 | -2725927.76 |
| G2 | -2725951.10 |
| G3 | -2725162.55 |
| G4 | -2725180.81 |

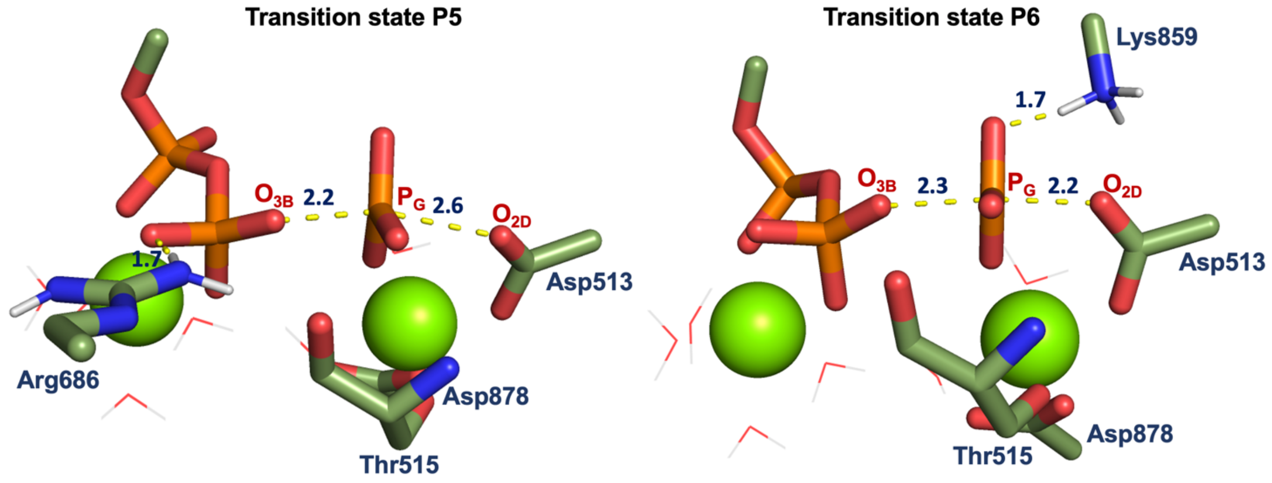


**Figure S8.** Transition state for systems P5 and P6. The transition state where Lys859 is missing is considerably affected (P5) while in system P6 the missing Arg686 does not influence the distances between the reacting nucleophile $O_{2D}$ and $P_G$ and $P_G$ and $O_{3B}$.
